# Early life adversity enhances prosocial helping towards a cagemate and reshapes brain-wide neural activity and network organization

**DOI:** 10.64898/2026.10.23.761382

**Authors:** Jocelyn M. Breton, Audrey Lynch, Janey Chiang, Samadhi Wijethunga, Isabelle Abdallah, Jeremiah Hickman-Maynard, Zara Poon, Inbal Ben-Ami Bartal, Heather Brenhouse

**Affiliations:** Smith College, Program in Neuroscience; Northeastern University, Department of Psychology; Tel Aviv University, School of Psychological Sciences

**Keywords:** Early Life Adversity, Maternal Separation, Prosocial Behavior, Social Motivation, c-Fos, Functional Connectivity

## Abstract

Early life adversity (ELA) is linked to changes in social behavior and emotional regulation and to increased risk for developing mood and conduct disorders. However, the effects of ELA on empathy and its associated behaviors, including prosocial helping, remain poorly understood. Animal models of ELA provide a tractable way to study how early rearing conditions impact prosocial behavior and underlying neural circuitry. In the current study, adult male and female rats reared under maternal separation (MS) or control conditions were tested in a helping behavior task in which they could release a trapped cagemate. Behavior was assessed across repeated testing sessions and brain-wide neural activity was quantified using c-Fos on the final test day. Contrary to our hypothesis, we found that MS-reared rats showed more consistent helping behavior, with shorter latencies to help on average, compared to control-reared animals. In addition, MS-reared rats displayed increased social play following release. At the neural level, MS-reared rats displayed more c-Fos expression across multiple brain regions, including prefrontal, amygdala, sensory, and striatal regions involved in motivation and social decision making. Network analyses further revealed altered functional connectivity in ELA-reared animals, with heightened connectivity in stress-related regions such as the lateral habenula. Together, these findings demonstrate that ELA impacts prosocial helping behavior and is associated with widespread changes in neural activity and network organization. This work provides insight into how early adversity shapes social motivation and the neural systems supporting prosocial behaviors.

## 1. Introduction

Early life experiences, including interactions with caregivers, critically shape socioemotional development and set the foundation for lifelong social behavior (Packard and Opendak, 2022; Wu et al., 2020). As a result, exposure to early life adversity (ELA), such as neglect or abuse, can lead to maladaptive social responding later in life (Tottenham, 2020). ELA also significantly increases the risk for developing mood and conduct disorders (Bomysoad and Francis, 2020; Heim et al., 2010; Heim and Nemeroff, 2001; Hunt et al., 2017; Kessler et al., 1997; Rytilä-Manninen et al., 2014; Spatz Widom et al., 2007), which are often characterized by changes in social engagement. A central component of social behavior is empathy – the ability to perceive and respond to the emotional state of others (Decety, 2015). Empathy involves concern for another’s wellbeing, which can motivate prosocial behaviors such as helping (Decety et al., 2016). These prosocial behaviors (actions that benefit another individual) are crucial for the establishment of positive relationships (Brown and Brown, 2015; Martela and Ryan, 2016), and enhance the wellbeing of both the actor as well as the recipient (Curry et al., 2018). Studies in humans suggest that ELA can impact empathy and empathy-related actions later in life, though evidence is mixed (Kahhale et al., 2025). For example, some groups have found that experiencing trauma early in life increases empathy (Benz et al., 2023; Greenberg et al., 2018) and prosocial actions (Lim et al., 2025, 2025; Vollhardt, 2009; Vollhardt and Staub, 2011). Yet, others have found that ELA blunts empathy and prosocial responding (Ardizzi et al., 2016; Bevilacqua et al., 2021; Cerqueira and Almeida, 2023; Fourie et al., 2019; Grimm et al., 2017; Huiping Zhang et al., 2024). These findings raise the question of when and how early adversity leads to heightened versus diminished empathy; indeed, the type of adversity experienced, and its timing may contribute to these varying findings (Chen et al., 2023; Kahhale et al., 2025). In addition, the neurobiological mechanisms by which early adversity shapes empathy and prosocial behavior remain poorly understood.

Animal models provide a powerful approach for controlling the early environment and for assessing how ELA impacts social behavior (Packard and Opendak, 2022; Parise et al., 2023). Notably, rats are sensitive to the affective states of conspecifics (Keysers et al., 2022) and demonstrate a variety of prosocial behaviors, including consolation of distressed individuals (Burkett et al., 2016), harm-aversion (Hernandez-Lallement et al., 2020), resource-sharing (Hernandez-Lallement et al., 2014), and helping or rescue behaviors (Bartal et al., 2011; De Waal and Preston, 2017; Decety et al., 2016; Gachomba et al., 2024), supporting their use for studying empathy-related processes. Though there is extensive work demonstrating how early adversity alters other domains of social behavior such as social approach, preference, and social memory ((Bolton et al., 2018; Parise et al., 2023; Raineki et al., 2012; Shin et al., 2018; Varlinskaya and Spear, 2008), relatively few studies have tested how ELA influences prosocial behaviors. Findings from these studies are mixed and dependent on both the type of adversity and behavioral assay. For example, while 24-hour maternal deprivation has been reported to have no effect on helping behavior (Kalamari et al., 2021), prolonged maternal separation has been linked to reduced prosocial responding across multiple paradigms (Joushi et al., 2022; Rombaut et al., 2023), highlighting the need for further work using comparable models and assays. Critically, though a small number of studies have begun to examine the effects of ELA on prosocial behaviors using animal models, mechanistic insight into these behavioral changes is lacking.

Prosocial behaviors such as helping are supported by coordinated activity across neural systems involved in emotional processing, motivation, and social decision-making (Luo, 2018; Wu and Hong, 2022). These processes engage a distributed set of regions, including areas classically associated with empathy such as the anterior cingulate cortex (ACC) and anterior insula (AI) (Bernhardt and Singer, 2012; Decety, 2015), as well as regions involved in reward, social evaluation, and memory, including the nucleus accumbens (NAc), prefrontal cortex (PFC), and hippocampus (Ben-Ami Bartal et al., 2021; Breton et al., 2022; La Greca et al., 2026; Wu and Hong, 2022). Converging evidence from humans and rodents suggests substantial overlap in these systems, supporting the existence of conserved neurobiological mechanisms underlying prosocial behavior (Decety et al., 2016). Together, these regions form an integrated “prosocial brain network” (Ben-Ami Bartal et al., 2021; Breton et al., 2022; Ruzal et al., 2026), within which coordinated activity is thought to support empathic responding and helping behavior. Notably, many of these same regions are highly sensitive to early life experience and are altered by ELA (Hanson et al., 2021; Malave et al., 2022; McLaughlin et al., 2019; Smith and Pollak, 2020), suggesting a potential pathway through which early adversity may shape prosocial behavior. However, despite evidence that ELA impacts these neural systems, it remains unclear how such alterations influence network-level activity during prosocial behavior.

To address these gaps, the present study examined how early life adversity shapes prosocial helping behavior and its underlying neural circuitry in rats. Specifically, we tested whether maternal separation (MS) alters helping behavior in adulthood and whether these behavioral changes are associated with differences in brain-wide neural activity and functional connectivity. Adult male and female rats reared under control or MS conditions were assessed using a helping behavior task (HBT), a well-established paradigm of targeted prosocial behavior (Bartal et al., 2011). Neural activity was then quantified across the brain to examine how ELA influences coordinated activity within circuits supporting prosocial responding. Importantly, this design allowed us to assess sex differences, addressing a critical and understudied factor in the field. Based on prior work demonstrating disruptions in social behavior following ELA (Parise et al., 2023), we hypothesized that ELA would impair helping behavior and reduce activity within neural circuits associated with prosocial behavior. By integrating a well-characterized model of early adversity with a validated measure of helping behavior and brain-wide analysis of neural activity, this study provides a framework for linking ELA-induced alterations in social behavior to underlying neural systems.

## 2. Methods

### 2.1 Animals and Housing

Sprague Dawley rats originally obtained from Charles River Laboratories (Wilmington MA) were bred in-house and maintained on a 12 h:12 h light-dark cycle (lights on: 0700-1900), in a temperature and humidity-controlled animal facility. All animals received *ad libitum* access to food and water and were housed in standard polycarbonate cages, with pine shaving bedding and a plexiglass tube for enrichment.

Litters were randomly assigned to either control (CON) or maternal separation (MS) conditions. Pups from 20 litters (10 CON, 10 MS) were included in the current experiments. Litters were culled to 12 pups on postnatal day (p) 1, with 6 males and 6 females per litter when possible. All rats were weaned and sex-segregated on p21, with most rats pair-housed with siblings of the same sex. Two cages (one CON male and one MS male) contained rats paired from different litters due to odd numbers. In total, 40 cages (80 animals) were used for the current experiment. A full description of all rats used, including their litter information, can be found in **Supplemental Table S1.**

All procedures were approved by Northeastern University’s Institutional Animal Care and Use Committee (IACUC) and were in accordance with the National Institute of Health’s Guide for the Care and Use of Laboratory Animals.

### 2.2 Maternal Separation (MS)

Maternal separation was conducted as previously described (Granata et al., 2022b; Lynch et al., 2026). Pups in litters assigned to MS conditions underwent daily separations from p2-p20. From p2-p10, each pup was placed in an individual plastic cup with a small amount of home cage bedding; the cup was placed in a circulating water bath (37C) for 3.5 hours. From p11-p20, each pup was placed in an individual mouse cage with bedding for 4 hours. Separations occurred at the same time each day (0930-1330), and dams remained in their home cage in the housing room during the separation period. CON and MS pups were weighed on p9 and p20. Control litters were also handled briefly on p12 and p15 to control for the minimal handling during maternal separation; otherwise, these litters were left undisturbed.

### 2.3 Experimental Design

This experiment followed a 2 (CON vs. MS) by 2 (Male vs. Female) design, with rearing condition and sex as independent variables. 10 cages of pair-housed rats were tested in each condition (CON Male, CON Female, MS Male, MS Female, n=10). After weaning at p21 animals were left undisturbed until they reached adulthood (>p65). Animals began the helping behavior test (HBT – described below), at p70-p75, on average. All testing was done in the light cycle.

### 2.4 Helping Behavior Test (HBT)

The helping behavior test (HBT) was performed as described previously (Ben-Ami Bartal et al., 2021; Breton et al., 2022). Prior to beginning the HBT, animals underwent five days of handling and habituation. On days 2-4 of that habituation, both cagemates were allowed 30 minutes to explore the empty arena (50 x 50 cm) together. On day 5, animals individually underwent a 15-minute open field task in the same arena. On all five of these days, animals were also tested for ‘boldness’, recording how long it took each animal to explore over the edge of an open homecage (Bartal et al., 2011; Hazani et al., 2025). The more ‘bold’ rat of the pair was assigned to be the ‘free’ rat for the helping test, while the less bold rat was assigned to the ‘trapped’ condition.

For the HBT, rats underwent 1 hour testing sessions over a 12-day period. On each day, the ‘trapped’ rat was placed inside a plexiglass restrainer, with a custom-made door that could be opened only by the ‘free’ rat on the outside. The door could be opened by pulling from the top or side or pushing up on the door with its snout. A successful door opening was counted if the free rat opened the door prior to the 40-minute mark. As in prior work, if the free rat did not open the door after 40 minutes, the door was propped open such that the trapped rat could easily push the door open from the inside with its snout. Both rats remained in the arena for the full hour. One week after the 12 days of testing, rats underwent three final test days. On the last day, the door was taped shut throughout the 60-minute session, such that it could not be opened; this allows all rats to have a similar experience on the final test day regardless of their history of helping and is intended to capture attempts to open the door rather than any social reward or play upon opening the door. All animals were perfused within 30 minutes of completing this final behavioral test (see details below). All sessions were video recorded with Logitech C922 Webcams using OBS studio software. Ethovision video tracking software (Noldus) was used to acquire and analyze all videos.

### 2.5 Behavioral Scoring and Analysis

Social interactions in the arena were scored both prior to the start of the HBT and throughout the HBT itself using Ethovision’s manual scoring: 30 minutes were scored on habituation day 2 and 10 minutes were scored on the first day of the HBT, day 6, and on the final test day in which animals could interact. Scoring was conducted as described previously (Breton et al., 2024), with two main categories of interactions: active interactions (including nose to nose and nose to body contact) and passive interactions (body to body contact without other movement). Several specific types of active interactions were also scored including play fighting (wrestling and pinning), anogenital sniffs and allogrooming.

### 2.6 Perfusions and Brain Extractions

Rats were euthanized within 90 minutes from the start of the final testing session, in order to capture expression of the immediate early gene c-Fos during early attempts to open (Lara Aparicio et al., 2022). Rats were anesthetized with CO2 and then transcardially perfused with ice cold 0.9% saline followed by freshly made 4% paraformaldehyde (PFA). Brains were post-fixed for a minimum of 18 hours in PFA, then sunk in 30% sucrose in 0.1 M PFB over several days. Brains were then stored at -80°C until they were ready to be sliced.

### 2.7 Histology and Immunohistochemistry (IHC)

Frozen brains were cryosectioned at 40 microns on a Leica cryostat (Leica 6800), starting around Bregma +4.70 and ending around Bregma -8.72 (Paxinos and Watson, 2006). Free-floating sections were stored in antifreeze in a 12-well plate, with every 12^th^ slice in the same well, following prior protocols (Ben-Ami Bartal et al., 2021). Samples were stored at -20°C prior to straining. IHC for the immediately early gene c-Fos was used as a proxy of neural activity during the final testing session. Tissue slices from one whole well (every 12^th^ slice) were stained for c-Fos, allowing for representation across much of the brain. Slices were first washed in tris-buffered saline (TBS) and blocked with 3% normal goat serum (NGS) in TBS with 0.3% Triton-X 100 for 1 hour at room temperature. Slices were then incubated overnight at 4°C with c-Fos primary antibody (1:1000, Abcam (ab190289) rabbit anti-cFos) in 0.3% TritonX-TBS containing 1% NGS. The next day, following rinses in TBS (3 x 5 minutes), sections were incubated for two hours in secondary antibody (1:500, Invitrogen (A11034) Alexa Fluor 488 goat anti-rabbit) in 0.3% TritonX-TBS with 1% NGS. Slices were rinsed in TBS again (3 x 5 minutes) and incubated for 10 minutes with 4’,6-diamidino-2-phenylindole (DAPI) (1:40,000 in 1xPBS), followed by additional rinses in TBS (4×5 minutes). A subset of tissue from the PFC only (between Bregma +4.7 to +1.6) was co-stained for the excitatory marker CaMKII following the same procedure and using the following antibodies (primaries: 1:1000, Abcam (ab190289) rabbit anti-cFos; 1:500 Cell Signaling Technology (#50049S) mouse anti-CaMKII; secondaries: 1:500 Jackson Immuno: Alexa Fluor 488 and 647). All slices were mounted on glass slides, coverslipped using 1,4-diazabicyclooctane (DABCO), dried overnight and stored until imaging.

### 2.8 Fluorescence microscopy and image analysis

All immunostained tissue was imaged at 10x using a wide field fluorescence microscope (Zeiss AxioImager M2) and processed using ZEN software (Zeiss). Regions of interest (250×250 micron squares) were placed across the brain as previously described (Ben-Ami Bartal et al., 2021; Breton et al., 2022), with several additional regions added (**Supplemental Figure S1**). A full list of the brain regions analyzed can be found in **Supplemental Table 2**. A custom written script in ImageJ/FIJI (Schindelin et al., 2012), was used to quantify c-Fos+ cells, with a set threshold for fluorescent intensity above background, and a threshold for cell size between 25-125 *μ*m^2^. CaMKII+ cells and co-staining with c-Fos was counted manually using ImageJ’s counting plugin. Any script-analyzed counts were also manually checked by multiple individuals blind to the animal’s sex and rearing condition. Inter-rater reliability was above 90%.

### 2.9 Statistical Tests

All data are presented as the mean +/-the standard error of the mean (SEM). Statistical analyses described below were performed using MATLAB, R, and GraphPad Prism. Outliers were a priori as values exceeding ± 2 standard deviations from the group mean and were excluded from analysis. In all tests, the alpha value was set to 0.05. Findings with p-values between 0.05 and 0.10 were considered trend-level. Effect sizes were calculated for all primary analyses and are reported where appropriate. Group sizes of 10 rats per condition were determined apriori using the G*Power tool, based on effect sizes reported in prior helping behavior studies (Ben-Ami Bartal et al., 2021; Breton et al., 2022), assuming alpha = 0.05 and power = 0.8 for between-group comparisons.

For most behavior analyses, two-way ANOVAs were used to assess main effects of rearing condition (CON vs ELA) and sex (males vs females). Repeated measures three-way ANOVAs were used when time was included as an additional variable. Mixed-effects analyses were used for datasets with missing values (for example, if an outlier was removed). A Kruskal–Wallis test was used to assess group differences in late-stage helping consistency (0–3 days of opening across the final three test sessions). A Fisher’s exact test was used to compare proportions of overall openers across groups. Planned post-hoc comparisons were run comparing ELA and CON animals within each sex for all measures. For c-Fos analysis, a three-way ANOVA was first run to assess main effects of sex, rearing, and brain region. As no main effect of sex was found, a two-way ANOVA was next run collapsing across sex and correcting for multiple comparisons using a Tukey’s post-hoc test. A False Discovery Rate (FDR) method was further used to correct for multiple comparisons following multiple t-tests on each brain region (Benjamini, 2006). Pearson’s correlations were also used to analyze relationships between behavioral metrics and between behavior and c-Fos measures. Additional statistical details can also be found within the Results section and within figure legends.

#### 2.9.1 Task partial least square (PLS) and network analyses

A Task PLS technique was used as previously described (Ben-Ami Bartal et al., 2021; Breton et al., 2022). To identify latent variables (LVs), permutation testing was performed using 500 iterations of resampling without replacement and shuffling of the test conditions. Following each resampling, the PLS was recalculated to determine whether effects of an LV were significantly different from noise. To assess salience (patterns that are maximally expressed in a given LV), bootstrap tests were performed 500 times using resampling with replacement while keeping subjects assigned to their group conditions. Brain regions with bootstrap ratios greater than 2.55 (denoted by a dashed line in the figure) were considered to significantly contribute to the pattern differentiating groups. For the PLS analysis, missing values were interpolated by using the average for that given group.

Network graphs based on c-Fos activity were also generated as previously described (Ben-Ami Bartal et al., 2021), visualizing the top 10% of correlations from a correlation matrix (using r values from pairwise Pearson correlations). Correlation values above that threshold were assigned a value of 1 and visualized as part of the network, with a line connecting the two brain regions in that correlation. Degree was calculated as the number of connections to a given node (brain region), while Betweenness was calculated as the number of connections moving through a node. The top 20% brain regions were identified for both Degree and Betweenness; regions that were in this top percentile for both were considered to be central hubs.

## 3. Results

### 3.1 ELA-reared rats helped release a trapped cagemate more than control-reared animals

Male and female rats reared under either early life adversity (ELA) or control (CON) conditions were tested in a helping behavior test (HBT) once they reached adulthood (n=10 per group; **Fig. 1A)**. Briefly, in this task rats are given the opportunity to learn to open a restrainer door, thereby releasing a trapped conspecific. This assay occurs over several weeks and rats are not trained in any way prior to testing (**Fig. 1B**). Prior work has demonstrated that control-reared adult male and female rats are motivated to help release trapped cagemates, with 50-60% learning to open consistently by the end of testing (Bartal et al., 2011; Ben-Ami Bartal et al., 2021; Hazani et al., 2025). Here, adult rats were tested with their trapped cagemates of the same rearing condition. Like controls, ELA-reared rats of both sexes learned to release a trapped cagemate, with many rats across all groups even learning to open the door on the first few days of testing (**Supplemental Figure S2**). As so many animals learned to open early on in testing, there was no difference in latency across the testing days (Time: F(13,459)=1.257, n.s. p=0.2358). We also found no main effects of sex or rearing condition on helping latency (sex: F(1,36)=0.05960, p=0.4451; rearing: (F(1,36)=2.601, p=0.1155). Although not statistically significant, ELA-reared rats exhibited faster latencies to help, on average, compared to controls (**Fig. 1C-D**). This same pattern was observed when averaging helping latencies across test days (F(1,36)=2.653, p=0.1120; **Fig. 1E**). Inspection of individual opening data across animals **(Fig. S2)** identified a subset of animals who opened inconsistently, contributing to high variability in latency measures. Thus, we next examined metrics of helping consistency, including the number of total days animals helped across the testing session. Here, we observed a significant main effect of rearing condition (F(1,36)=4.144, p=0.0492, ηp^2^=0.103; **Fig. 1F**), with ELA-reared animals helping on more days relative to controls. No significant effect of sex and no interaction between sex and rearing were observed. To further assess stability of helping behavior, we analyzed helping across the final three test days. A Kruskal–Wallis test revealed a significant effect of rearing condition on late-stage helping consistency (χ²(1) = 5.39, p = 0.020), with ELA-reared rats showing a shift toward higher consistency scores (**Fig. 1G)**. The effect size was moderate-to-large (η²H = 0.116), indicating a meaningful shift in the distribution of helping behavior across the final test days. Finally, we compared the proportion of animals classified as fully consistent ‘openers’ (defined as opening on all three of the last possible days (days 12-14). Here, ELA reared rats showed a trend towards a greater proportion of consistent helpers (Fisher’s exact test: p=0.068), with 5/10 CON and 8/10 ELA females ultimately becoming ‘openers’, and 3/10 CON and 8/10 ELA males, respectively (**Fig. 1H**).

**Figure 1.**
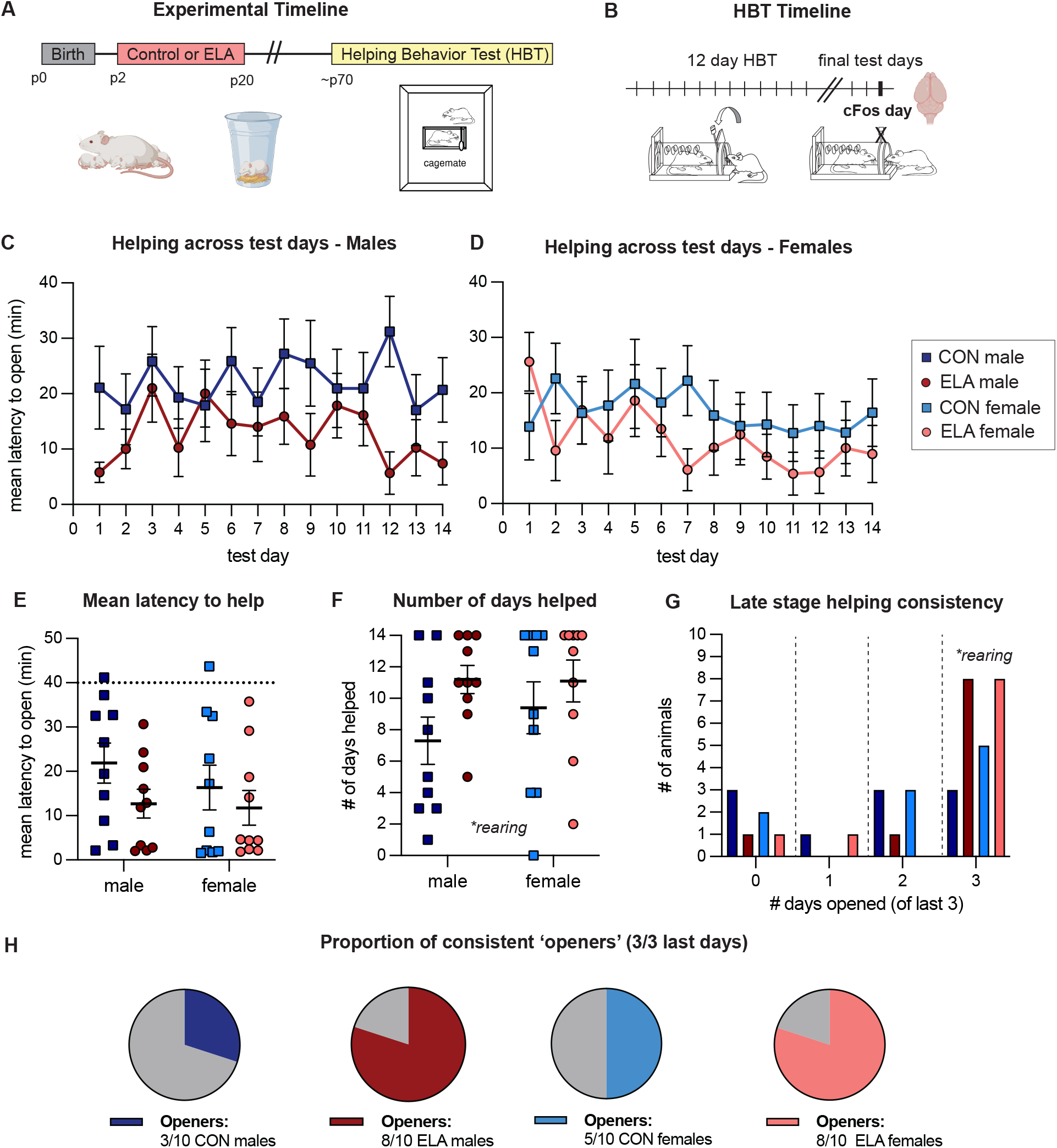
ELA influenced helping behavior in adult rats. A) Experimental Timeline. Animals were reared under control (CON) or early life adversity (ELA) conditions from postnatal day (p) 2-p20. All animals started the helping behavior test (HBT) in adulthood. B) Animals underwent 12 days of the HBT followed by a one-week break. There were three final test days; on the last day the door was locked shut such that the free rat could not open it. Brains were collected in a time-locked manner for subsequent analysis. C-D: Helping behavior across the test days in male (C) and female (D) rats. Shown is the average latency to open the door (in minutes) per group. E) The mean latency to open the door averaged across all test days. The dashed line at 40 minutes represents when the door was halfway opened by the experimenter. F) The overall number of days with successful helping by group. G) Late stage helping consistency across the last testing days where animals were able to help (days 12-14). Shown is the number of animals per group who helped on either 1, 2 or 3 of these three testing days. H) The overall proportion of rats that became fully consistent ‘openers’, defined as opening on 3/3 of these last days 12-14. n=10 per group.

Overall, these unexpected findings demonstrate that ELA-rearing in the form of maternal separation does not impair prosocial motivation towards a trapped cagemate. Instead, ELA-reared rats exhibited increased helping behavior, characterized by a greater number of helping days and a shift toward more consistent late-stage helping behavior.

### 3.2 ELA-reared rats spend more time interacting with their trapped cagemate on the final day of testing

On the final day of testing, the restrainer door was latched shut for the full duration, such that all rats similarly experienced exposure to their trapped cagemate, despite their different histories in door opening. Movement patterns, such as time spent around the restrainer, (**Fig. 2A-B**) were analyzed to assess motivation to interact with their cagemate. On this final test day (day 15), ELA rats spent more time around the restrainer than controls (main effect of rearing: F(1,35)=7.909, p=0.008, ηp^2^=0.184; **Fig. 2C**). Though there was no main effect of sex, there was a trend for an interaction of sex and rearing (F(1,35)=3.228, p=0.081), and a-priori planned posthoc comparisons within each sex found that ELA males, but not females, spent more time interacting with their trapped cagemate compared to their control counterparts (Sidak: p=0.0044; **Fig. 2C**). The same effect was observed for the number of entries into the restrainer zone (**Supplemental Figure 3A**). Importantly, there were no significant differences in total distance moved (**Fig. 2D**) or velocity (**Fig. S3B**) across groups, indicating that ELA-reared animals were not simply more active overall, but instead, specifically differed in where they chose to spend time. The distance animals moved within the zone around the restrainer was also significantly correlated with the animal’s latency to help across all test days (Pearson’s correlation, r=-0.6427, p<0.0001; **Fig. 2E)**; animals that moved more around their trapped cagemate were faster to help, on average, on prior days. The percent of time spent around the restrainer and the number of entries into the restrainer zone were similarly correlated with helping latency, as was the velocity around the restrainer (**Supplemental Figure 3C-E**). However, correlations with helping latency were not observed with the total distance moved around the edges of the arena (**Fig. 2F),** or with velocity moved around the edges (**Fig. S3F**), indicating that, on this final test day, movement specifically directed towards their trapped cagemate was reflective of prior motivation to help. Combined, these data mirror findings across the HBT testing sessions, with ELA-reared rats showing heightened motivation to engage with and help their trapped cagemate.

**Figure 2.**
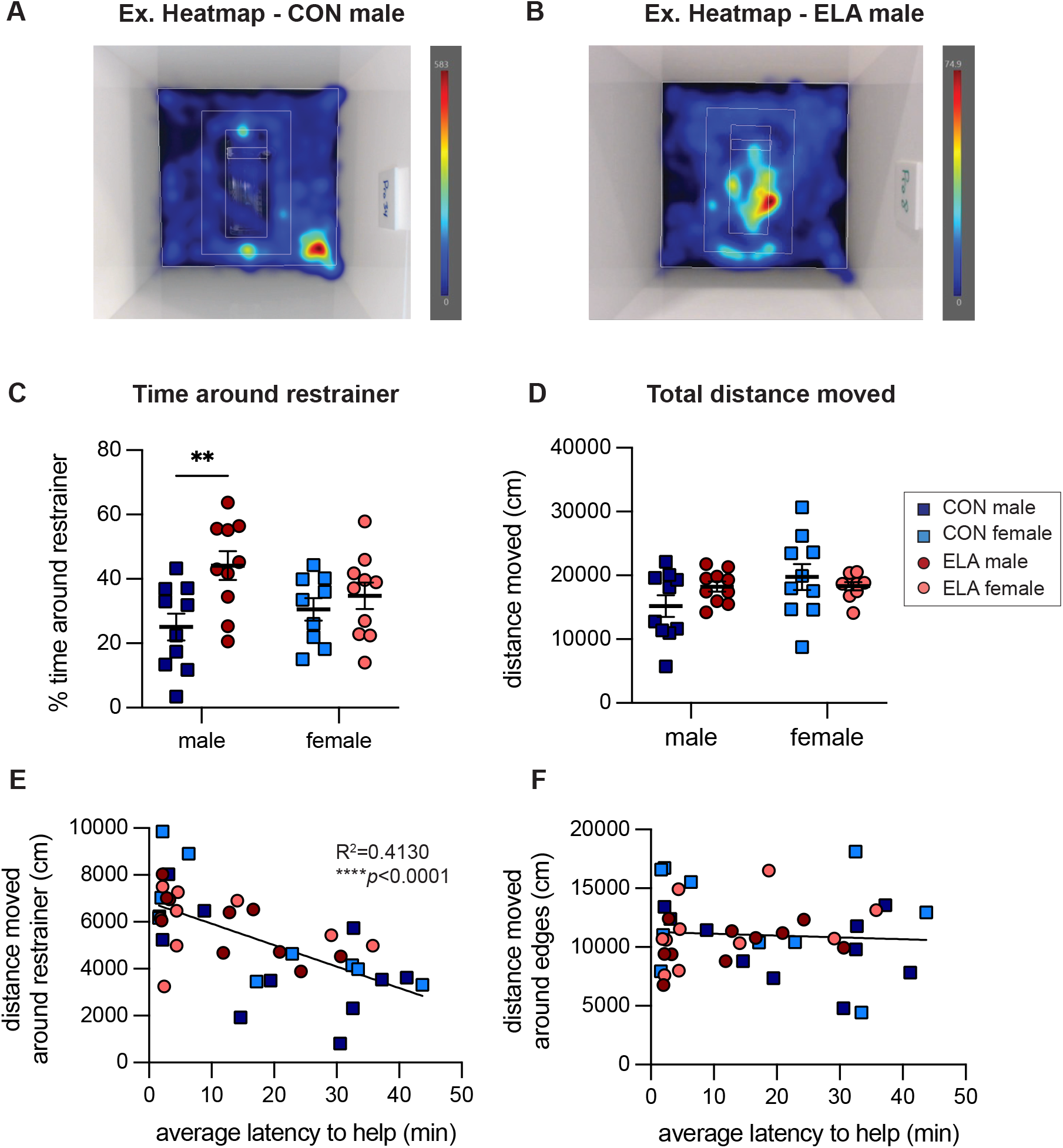
ELA-reared rats spent more time with their trapped cagemate on the final test day. (A-B): Representative heatmaps of movement patterns from a control-male (A) and ELA-male (B) on the final test day (day 15). Zones are shown including the zone around the restrainer. C) Percent of time spent in the zone around the restrainer across the 1-hour session. D) Total distance traveled on the final session. (E-F): The distance moved within the zone around the restrainer was significantly correlated with the animal’s latency to help on prior days (E) but was not correlated with distance moved around the edges of the arena (F). n=10 per group.

### 3.3 ELA-reared rats show increased play fighting behavior upon door opening

Social interactions upon door opening may provide insight into the motivational state of rats tested in the HBT (Breton et al., 2022; Hazani et al., 2025). Here, we analyzed social interactions throughout the HBT paradigm, including at the start of testing, mid-way through and at the end of the testing period (**Fig. 3A**). Though the total duration and number of social interactions did not differ across groups (**Fig. 3B, Supplemental Fig 4A**), the types of interactions varied depending on sex and rearing condition. Specifically, the total duration and number of play fights differed based on rearing (duration: (F(1,36)=5.938, p=0.0199, ηp^2^ = 0.1416; **Fig. 3C**; number: F(1, 36)=5.845, p=0.0208, ηp^2^ =0.1397; **Fig. S4B**), with no significant effect of sex and no interaction between sex and rearing. In contrast, play fights were rarely observed in any condition during the habituation days prior to beginning the helping behavior test (**Fig S4C**), indicating that play fights were unique to the animal’s state upon releasing the trapped rat. While play fighting during the HBT differed based on rearing, anogential sniffs, by contrast, showed a main effect of sex (F(1,36)=12.81, p=0.0010, ηp^2^ =0.2624; **Fig. 3D**), but no effect of rearing nor interaction between sex and rearing. Here, females sniffed more, on average, compared to males (CON F: 32 ± 3.6; CON M: 22 ± 4.2; ELA F: 34.3 ± 5.2; ELA M: 16 ± 2.2). Lastly, allogrooming duration differed based on sex (F(1,36)=7.949, p=0.008, ηp^2^= 0.1895; **Fig. 3E**), but there was no main effect of rearing condition nor interaction between the two. This pattern in allogrooming was mirrored when examining the number of allogrooming events (**Fig S4D**). Three-way repeated measures ANOVAs looking across the testing days also revealed similar main effects of rearing condition and sex for each social behavior (**Fig. S4E-G**). Though there were significant effects of test day for most social behaviors, no interactions between sex and rearing and test day were observed, indicating differences across groups were present even on the early test days (for full statistics, see **Supplemental Table 3**). Overall, these results highlight that ELA-reared rats of both sexes engaged in more rough and tumble play with their cagemate in the 10 minutes upon their release, perhaps pointing to differential motivational states for helping. Importantly, early rearing environments changed the type of social interactions observed rather than changing the overall amount of social interaction.

**Figure 3.**
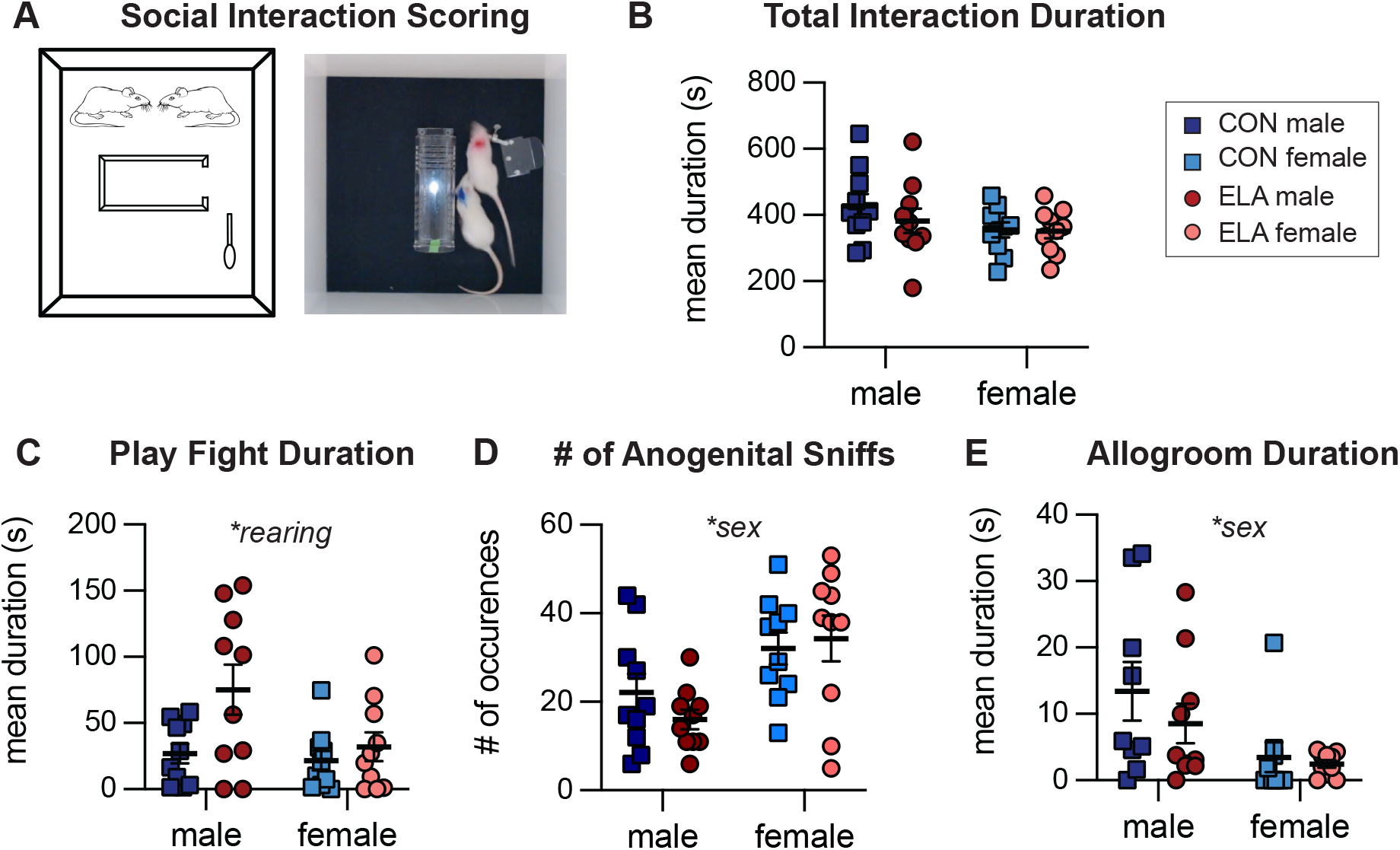
ELA-reared rats showed increased play fighting upon release of their trapped cagemate. A) Diagram and representative image of social interaction between two cagemates upon opening the restrainer door. B) The total duration of group social interactions across all social interaction scoring. C) Play fight duration D) The number anogenital sniffs and E) Allogrooming duration. n=10 per group.

### 3.4 ELA-reared rats of both sexes show higher levels of brain-wide c-Fos on the final day of testing compared to controls

Brain-wide activation patterns were compared between ELA and CON reared rats on the final day of testing, using the immediate early gene c-Fos as a proxy of neural activity (**Fig. 4A-B, Fig. S1)** (Chung, 2015). All rats had the same experience on the final day of testing with the restrainer door latched shut (day 15), allowing us to measure markers of neural activity in response to a trapped cagemate; importantly, this reflects engagement with the trapped rat rather than social interactions and/or social reward upon opening. This experimental design was especially important, as most rats had learned to open within minutes of being placed in the arena and we would otherwise have captured signals representing social interactions after release. Further, this strategy has proven useful in the past for distinguishing neural signals across groups with different histories of helping (Ben-Ami Bartal et al., 2021; Breton et al., 2022).

**Figure 4.**
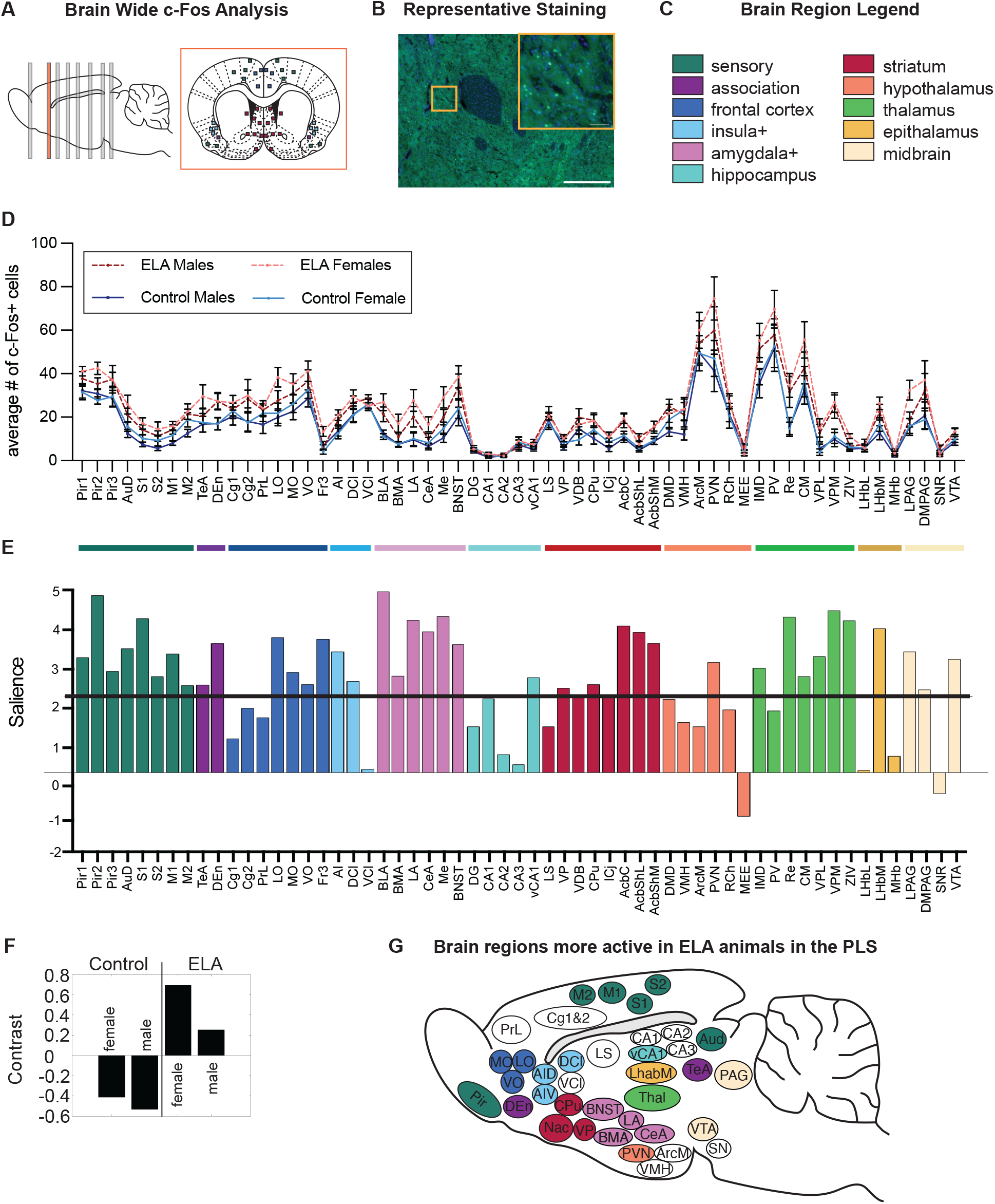
ELA-reared rats have higher levels of neural activity compared to controls. A) Diagram showing locations of brain slices used for c-Fos analysis and a representative section from A.P. 1.56. B) Representative c-Fos staining from the nucleus accumbens core. Scale bar is 500μm on the full image, and 50μm on the zoomed in square. C) Legend of brain region categories coded by color, used throughout the manuscript. D) Number of c-Fos+ cells per region (mean ± SEM). E) A salience plot (representing z-scores of bootstrapped tests), with regions crossing the black threshold line significantly (p<0.01) contributing to the contrast shown in (F). The directionality of the bars corresponds with the direction in the contrast graph below. G) A sagittal section with brain regions that crossed the salience threshold in (E) colored.

Several statistical methods were used to compare whole-brain neural activity across conditions. First, a mixed effect analysis was used to compare effects of rearing condition and sex across all brain regions (**Fig. 4C-D)**. A main effect of brain region was found, as expected (F(58,2086)=66.60, p<0.0001; ηp^2^= 0.6128), as was a main effect of rearing condition (F(1,36)=10.46, p=0.0026; ηp^2^= 0.195) and an interaction between brain region and rearing condition (F(1,36)=2.468, p<0.0001; ηp^2^= 0.045). However, no primary effect of sex was found, indicating male and females in each condition displayed a similar pattern of neural activity. As no main effect of sex was identified, brain data from both sexes were combined to better power analyses. A 2-way ANOVA with sexes combined again identified a main effect of brain region (F(66,2496)=57.92, p<0.0001, ηp^2^= 0.605), rearing (F(1,38) = 8.941, p=0.0023; ηp^2^= 0.191) and a brain by rearing interaction (F(66,2496)=1.695, p=0.0005, ηp^2^= 0.043; **Figure S5**). Interestingly, ELA animals showed elevated levels of c-Fos across much of the brain (**Fig S5**). A Tukey’s post hoc test for multiple comparisons was run next, comparing ELA and control rats in each brain region (**Supplemental Table 4).** Many brain regions showed significantly more c-Fos+ cells in ELA compared to control rats, including in sensory, prefrontal, amygdala and reward regions. As a secondary statistical approach, multiple unpaired t-tests were run for each brain region, followed by a false discovery rate (FDR) method used to correct multiple comparisons (Benjamini, 2006). A similar pattern was observed in almost all brain regions using these two methods (51/59 regions; **Table S4**).

Lastly, a multivariate Partial Least Square (PLS) analysis was used to identify patterns of brain activity that best differentiated the experimental conditions. PLS is an especially useful tool for brain-wide analyses as no assumptions are made and there is no need to correct for multiple comparisons; rather, the analysis aims to maximize the contrast between groups. Using data from all four groups, a primary significant latent variable (p<0.001) emerged associated with rearing condition, with little differences across sexes (**Fig 4. E-G**). A multitude of brain regions (39/59) significantly contributed to this contrast, as shown by brain regions that cross the salience threshold (**Fig. 4E; Table S4**), with some regions playing a bigger role in distinguishing between conditions, including the basolateral amygdala (BLA) and other subregions of the amygdala (shown in pink), regions within the nucleus accumbens (in red), and several sensory, prefrontal and thalamic nuclei (in dark green, dark blue and light green – respectively; **Fig. 4E**).

Similar effects were observed in 76% of cases (45/59 regions) across all three statistical methods, providing validation and support for use of these methodologies. Importantly, 28 brain regions showed statistically significant differences between control and ELA conditions across all three statistical methods employed (**Table S4**); these regions were the ones that contributed most to the contrast within the PLS and included the amygdala and nucleus accumbens subregions. Overall, these data suggest that ELA males and females have heightened neural activity in many regions of the brain on the final day of testing relative to control animals.

### 3.5 Neural activity in frontal cortical regions is correlated with performance in the helping behavior test

We next assessed whether brain wide activity markers measured via c-Fos were related to animals’ performance in the HBT. Here, correlation analyses were utilized to assess if c-Fos levels on the final day of testing were correlated with the average helping latency across prior days of testing (**Fig. 5A**). Five brain regions were significantly correlated with helping latency: the prelimbic cortex (PrL), lateral and ventral orbitofrontal cortices (LO and VO), the CA1 region of the hippocampus, and the median eminence (MEE) of the hypothalamus (**Fig. 5B-F)**. In these regions, higher levels of c-Fos were associated with faster helping latencies on average. It was notable that three of these regions were in the frontal cortex, especially as the PrL and regions of the orbitofrontal cortex (OFC) have been previously associated with conditions of helping (Ben-Ami Bartal et al., 2021; Breton et al., 2022).

**Figure 5.**
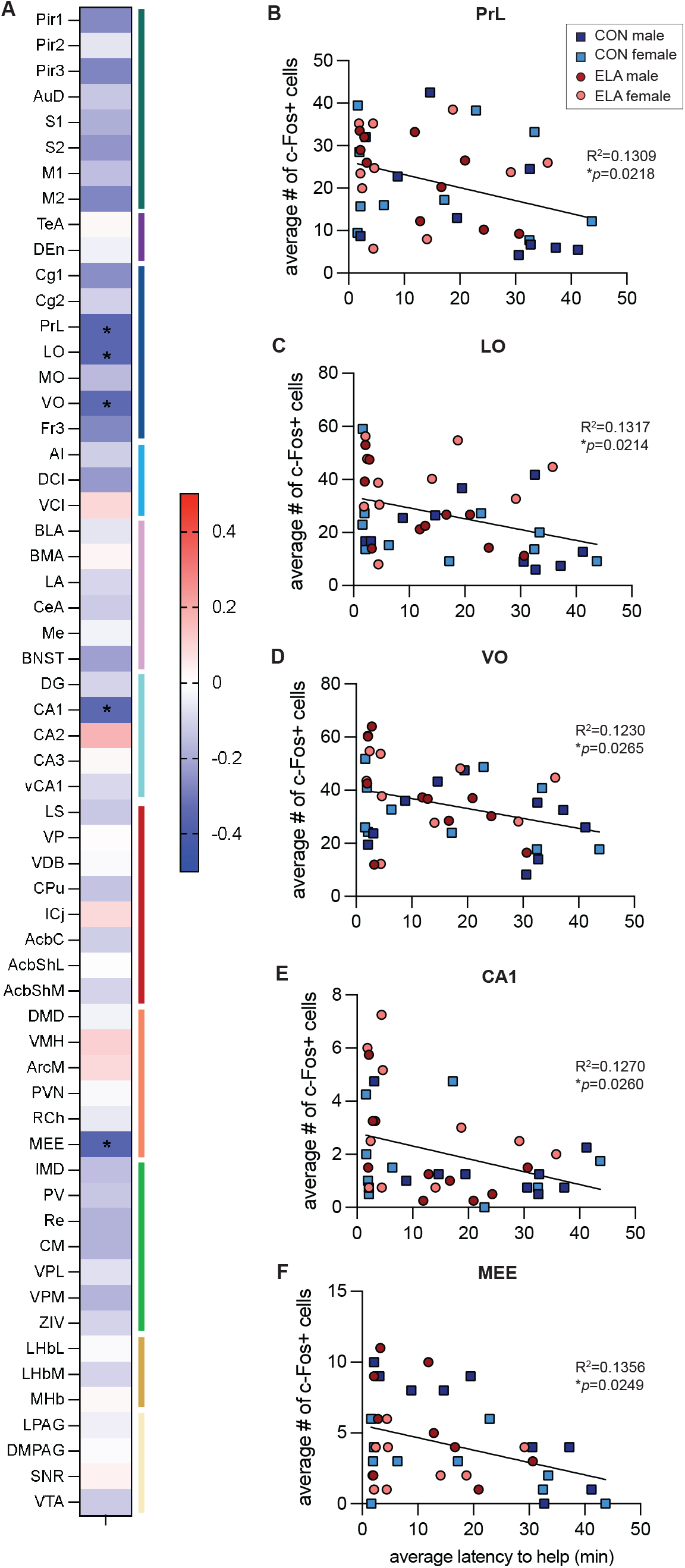
c-Fos levels in three prefrontal cortex regions were correlated with helping behavior. A) Heatmap of Pearsons’s correlations (r-values) showing correlations of c-Fos levels in each brain region with the average latency to help during the HBT. Red indicates positive correlations, while blue indicates negative correlations (more c-Fos associated with faster latencies to help). Statistically significant regions are denoted with asterisks. (B-E): Correlation graphs of the five significant brain regions: the PrL (B), LO (C), VO (D), CA1 (E) and MEE (F).

The PFC contains a large population of excitatory, glutamatergic neurons; within the PrL and OFC, these cells are thought to help regulate goal-directed and social-decision making behaviors (Klein-Flügge et al., 2022; Mack et al., 2024; Woon et al., 2020). Thus, here, we sought to test whether these behaviorally relevant c-Fos signals reflected differential recruitment of excitatory projection neurons within the PFC. Co-staining tissue for c-Fos and the excitatory marker CaMKII (Liu and Jones, 1996) revealed that roughly 50% of c-Fos+ cells in all PFC regions were CaMKII+, however no group differences were observed based on sex or rearing condition (**Supplemental Figure S6**). These data suggest that ELA does not lead to a global increase in excitatory drive within the PFC, and thus, bulk excitatory activity likely does not drive differences in helping behavior per se.

### 3.6 Network analyses show different patterns of functional connectivity in ELA and control-reared animals

As higher levels of c-Fos markers were observed across much of the brain in ELA animals, spanning multiple brain networks with various functions, we next asked whether brain-wide correlations of activity differed in ELA compared to control-reared animals. For example, it is possible that neural activity is simply heightened in ELA animals, but that metrics of functional connectivity are similar to control-reared animals, indicating that early life experience might impact salience but not change how regions are communicating with one another. In contrast, differences in network connectivity might indicate that early adversity impacts communication between regions when exposed to a trapped conspecific. To test these hypotheses, we used a ‘functional connectivity’ mapping approach, as previously described (Ben-Ami Bartal et al., 2021; Breton et al., 2022). Here, we first looked at correlations in all forty animals to see what emerged as overarching patterns in brain activity before looking at how this activity shifted in control vs. ELA reared animals. Using all animals, we saw three major clusters emerge (**Supplemental Figure S7**). The largest cluster contained brain regions that have been previously shown to be associated with prosocial behavior in this task (Ben-Ami Bartal et al., 2021), including sensory regions, insular cortex regions, prefrontal cortex regions including the OFC, as well as the nucleus accumbens shell – thus, we termed this cluster the “prosocial” cluster. The second cluster contained amygdala regions such as the BLA, central amygdala (CeA), lateral amygdala (LA) and medial amygdala (Me), as well as the bed nucleus of the stria terminalis (BNST), paraventricular nucleus of the hypothalamus (PVN), periaqueductal grey (PAG) and several thalamic subregions; many of these regions are canonically involved in stress and arousal ((Davis et al., 2010; Herman and Cullinan, 1997; McGaugh, 2004; Roozendaal et al., 2009; Hui Zhang et al., 2024) and thus we termed this the “stress/arousal” cluster. The final cluster consisted of areas associated with movement such as the caudate and putamen (CPu) and nucleus accumbens core (AcbC) (Groenewegen, 2003; Nicola, 2007), as well as socially motivated regions like the lateral septum (LS), the ventromedial hypothalamus (VMH) and the basomedial amygdala (BMA) (Ferrara et al., 2023; Menon et al., 2022; Modi and Sahin, 2019), along with several hypothalamus regions; we termed this cluster “social vigor” given those combined functions.

We next looked at network metrics for control (**Fig. 6A-E**) and ELA-reared (**Fig. 6F-J**) animals separately. Correlation matrices differed qualitatively across conditions; controls had a more coherent network, with strong correlations across much of the brain, while in contrast, ELA-reared animals had more discrete clusters. We then visualized the top 10% of correlations in each rearing condition as network maps, and color-coded brain regions according to the three primary clusters based on the entire group (**Fig. S7**). For controls, high connectivity was observed across all three clusters, with the lateral septum (LS) emerging as the largest hub (**Fig. 6C**), with both the highest number of connections (degree; **Fig. 6D**) and number of connections going through it (betweenness; **Fig. 6E**). For the ELA animals, the three clusters that emerged mapped well onto the original three based on the full group, with “prosocial” regions in purple isolated in their own cluster. The medial subregion (LhbM) of the lateral habenula (Lhb), a region associated with emotional integration and aversive signaling (Lecca et al., 2014; Zheng et al., 2024), emerged as the largest hub (**Fig. 6H**), with the highest degree (**Fig. 6I**) and betweenness scores (**Fig. 6J**). The betweenness score for the LbhM in the ELA group was also three times as great as that observed in controls. In addition, the LhbM appeared to be a connector between the rest of the brain and these prosocial- and sensory-related regions for ELA animals. Altogether, these results suggest that, upon exposure to their trapped cagemate, ELA animals had more functional connectivity between stress-related regions like the lateral habenula and prosocial and goal directed regions like the orbitofrontal cortex.

**Figure 6.**
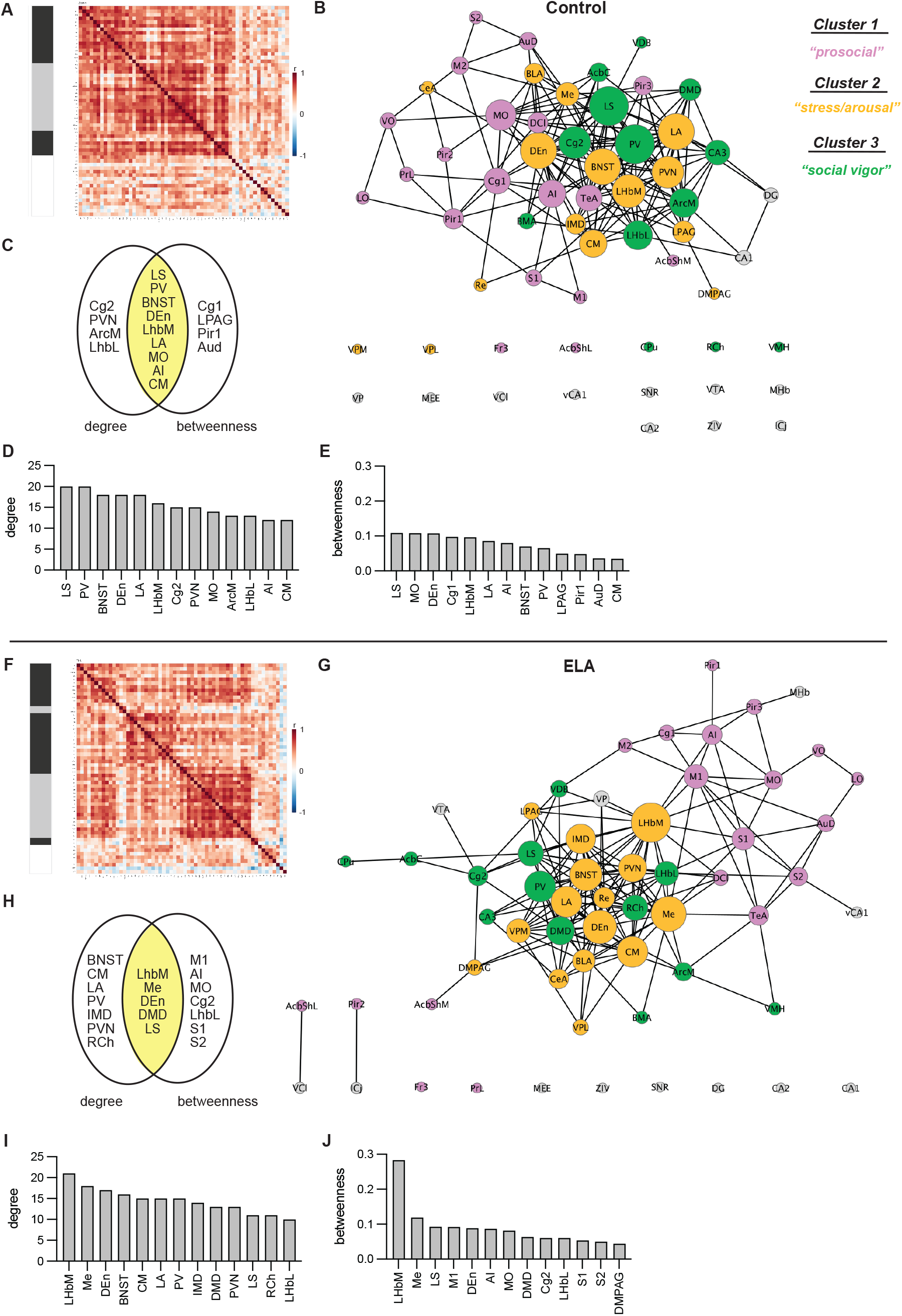
Different patterns of functional connectivity were observed in ELA compared to control rats. Correlation matrices (A,D), network maps (B, E) and central hubs (C-E, H-J) for control-reared rats (A-E) and ELA-reared rats (F-J). Networks are color coded based on the three main clusters that emerged from all animals – see supplemental figure S7. The “prosocial” cluster is in pink, the “stress/arousal” cluster is in orange, and the “social vigor” cluster is in green.

## 4. Discussion

In the current study, we tested how ELA, in the form of MS, influenced prosocial helping behavior and brain activity in adult male and female rats. ELA-reared rats showed increased helping toward a trapped cagemate relative to controls and exhibited increased social play behavior with the liberated animal. Neural activity during exposure to a trapped conspecific was elevated in ELA-reared rats of both sexes, especially in amygdala, sensory, and striatal regions. Functional connectivity analyses further revealed more discrete networks in ELA-reared rats, with the lateral habenula emerging as a key hub linking stress-related and prosocial circuits. Together, these findings suggest that ELA can reshape social motivation and reorganize connectivity across social brain networks. These results advance our knowledge of how early life experiences influence empathy-related behaviors and expand our understanding of the neural systems supporting prosocial helping.

### 4.1 Behavioral effects of ELA on prosocial helping

Our data add to a small but growing body of work investigating how ELA impacts empathy-related behavior in rodents. Multiple experimental paradigms have been used to test a variety of rodent prosocial behavior, including rescue-behavior, consolation, harm aversion, and reward-based prosocial choices (Gachomba et al., 2024). Our study is among the first to examine how ELA impacts rescue behavior. No other groups have tested the effects of prolonged MS on helping; however, Kalamari et al (2021) tested the impact of a single 24-hour maternal deprivation (MD) on rescue behavior using an operant based task (Kalamari et al., 2021). In contrast to our findings here, they found that MD did not influence motivation to help release a trapped cagemate. Thus, it is possible that different types of ELA (single 24 hr vs 3+ hrs across 20 days) lead to different outcomes in prosocial motivation. Furthermore, the liberation task used by Kalamari and colleagues relied on lever pressing, with extensive training prior to prosocial testing, while the current task did not shape or train release behavior in advance. Future work should consider the impact of prior training on behavioral performance, especially if training might minimize or mask differences across groups.

Differences in helping behavior may also be driven by changes in the caregiving environment. MS significantly alters maternal care in rats, with changes in maternal behaviors like licking and grooming as well as changes in the predictability of care (Baracz et al., 2020; Granata et al., 2022a; Rombaut et al., 2023). In a study by Asadi et al (2021) adult male rats who had received higher levels of maternal care showed an increased willingness to open a door to free a distressed, soaked conspecific (Asadi et al., 2021). Maternal care was not directly measured in the current study; however, this would be an interesting metric for future studies to analyze. For example, future work could test whether maternal behavior during ELA can predict future helping behavior.

Stress contagion is an important component in driving prosocial helping (Bartal et al., 2016). Of relevance, allogrooming distressed conspecifics is a form of prosocial behavior that depends on stress contagion and that has been found to be altered following ELA. In particular, both MS and limited bedding ELA paradigms lead to reduced consolation of stressed cagemates (Breton et al., 2024; Rombaut et al., 2023). In the current study, very little allogrooming was observed upon release, with no differences across rearing conditions. Direct helping very likely elicits a reward response upon release (Ben-Ami Bartal et al., 2021; Ruzal et al., 2026) and may rapidly mitigate stress levels of the trapped rat, minimizing a need for further consolation or prosocial actions. Future studies could aim to assess stress response levels both pre and post helping in the free and trapped animals, providing insight into the role of emotional contagion in helping behavior and its interactions with ELA. Altogether, there are a number of variables to consider that may impact prosocial motivation, including the type of adversity experienced and its timing, the specific form of prosocial behavior measured, and whether vicarious stress is experienced.

### 4.2 Sex differences in prosocial behavior

Biological sex is an additional factor to consider when assessing prosocial motivation, especially given work demonstrating sex differences in social behavior following ELA (Kundakovic et al., 2013). Of note, many of the aforementioned studies testing how ELA impacts prosocial behaviors only analyzed behavior in male rodents. Our study is the first to test for sex-differences in prosocial helping behavior following ELA. We found ELA led to increased helping in both male and female rats, with no main effect of sex on helping latencies. However, on the final day of testing, ELA-male rats spent a greater amount of time near their trapped cagemate compared to controls, while females of both rearing conditions showed similar behavior, indicating that sex does indeed interact with early life experiences to influence social behavior.

Past studies have demonstrated that females animals may be more prosocial than males (Avital et al., 2016; Bartal et al., 2011; Misiołek et al., 2023). Here, though we found that a greater proportion of control females helped (6/10) relative to control males (3/10), and that females tended to help with faster latencies than males on average, these differences were not statistically significant. A sex difference was observed, however, when examining social behaviors upon door opening. Specifically, in both rearing conditions, female rats showed greater anogenital sniffing following release. Importantly, stress signals are emitted from the anogenital region and are necessary for the social transmission of stress (Lee et al., 2021; Sterley et al., 2018). One possibility is thus that females are emitting and/or detecting more stress signals from their trapped conspecifics, and these signals may play a role in their motivation to help. Future studies could assess stress transmission in both sexes to test this hypothesis. Another possibility is that the same behavioral outcome (helping) is observed in males and females, yet is driven by different motivations (Bath, 2020). Future work should continue to include sex as a biological variable in order to understand the full extent of ELA’s impact on social behavior.

### 4.3 Social interactions upon prosocial helping

We examined social behavior between cagemates upon door opening at several points throughout the testing period. Interestingly, ELA-reared rats of both sexes showed increased play fighting behavior immediately upon release. This finding may provide insight into the animals’ motivation for helping; one possibility is that social reward upon helping is heightened in ELA conditions. It would be informative for future work to test therefore whether ELA-reared rats would be equally motivated to help if their cagemate were released into a separate compartment, without the possibility for direct social interaction. Indeed, some have found that helping behavior diminishes when social contact is removed (Kalamari et al., 2021), though others have reported that social contact is not necessary for helping to occur (Bartal et al., 2011; Ruzal et al., 2026). In general, MS can increase play fighting in peri-adolescent male rats (Lundberg et al., 2017; Veenema and Neumann, 2009), though this same effect has not been observed in females (Lundberg et al., 2017). It was thus surprising to see increased play fighting in both males and females, and further, in adulthood, when play fighting is often reduced relative to adolescents (Panksepp, 1981; Pellis and Pellis, 1990; Varlinskaya and Spear, 2008). Importantly, play fighting behavior was rarely observed in the home cage prior to testing, with no differences in baseline play behavior across all groups, indicating that this change in social behavior emerged specifically after release of the trapped rat. Understanding the factors that influence the motivation to help and how they interact with ELA will be an important area for continued study.

### 4.4 Neural activity marker changes following early adversity

Prosocial helping behavior involves the activation of a widespread network of brain regions, including sensory, prefrontal, striatal, and thalamic systems (Decety, 2015; Li et al., 2025; Luo, 2018; Meyza et al., 2017; Walsh et al., 2023). The overall patterns of c-Fos we observed here replicated those identified previously in animals with a history of helping (Ben-Ami Bartal et al., 2021; Breton et al., 2022). We found that ELA-reared males and females had elevated c-Fos levels across many regions of the brain relative to controls. One possibility is that this increase in c-Fos following ELA may be indicative of heightened sensitivity to their trapped conspecific; the specificity of neural activity marker increases should be confirmed in future work. Indeed, a recent meta-analysis found that ELA can drive increases in immediate early gene expression at rest across multiple brain regions (Schuler et al., 2022). Yet, differences in c-Fos levels between ELA and control-reared animals were dampened following exposure to acute stress (Schuler et al., 2022). In the present study, a partial least-squared (PLS) analysis allowed us to determine which regions most significantly contributed to group differences between ELA and control animals.

Of note, all regions in the amygdala displayed strong, statistically significant salience scores. The amygdala, especially the MeA, is well known for its role in social behaviors such as social reward and social approach (Li et al., 2017; Raam and Hong, 2021), as well as prosocial actions like comforting behaviors (Qu et al., 2024; Wu et al., 2021). Further, changes in MeA neural activity have been identified following ELA (Hu et al., 2021; Mañas-Ojeda et al., 2025), highlighting the MeA as a possible mediator underlying behavioral changes observed in ELA and control animals. The BLA is another key subregion that was significantly more active in ELA-reared animals on the final day of testing. The BLA processes emotional information, including auditory cues from other individuals (Ghasemahmad et al., 2024; Naumann and Kanwal, 2011), and it plays a role in promoting altruistic choices over selfish ones (Scheggia et al., 2022). Early adversity also highly impacts the development of the BLA, including connectivity with the prefrontal cortex (PFC) (Gee et al., 2013; Honeycutt et al., 2020). How and if ELA-driven changes in the amygdala are related to changes in motivated behavior is an important area for ongoing study.

Sensory regions were also significantly more active in ELA-reared rats relative to controls. The helping behavior task involves multiple sensory cues, including auditory and olfactory signals; such cues are inherently linked with affective processing in both rodents and in humans (Contestabile et al., 2021; Satpute et al., 2015; Zhang et al., 2018). Elevated c-Fos, especially in the piriform cortex, may indicate that ELA-reared rats show heightened sensitivity to olfactory cues released by their trapped cagemate. A recent study by Ruzal et al (2026) identified specific recruitment of oxytocin receptor positive (Oxtr+) cells in the sensory cortex following helping behavior (Ruzal et al., 2026). Future work could assess where differences in this specific cell type are observed in ELA rats, providing added insight into observed differences in prosocial behavior.

Notably, within the striatum, all subregions of the NAc significantly contributed to the contrast between groups. The NAc is well known for its role in reward and motivated behaviors (Robbins and Everitt, 1996; Salamone, 1994), and prior work has identified increased NAc activity in conditions of helping (Ben-Ami Bartal et al., 2021). Increases in NAc c-Fos in ELA compared to control animals may therefore be indicative of heightened social motivation. However, the exact role of the NAc in prosocial helping behavior is still under investigation. In prior work, c-Fos markers in the NAc were correlated with door opening behavior (Ben-Ami Bartal et al., 2021). Here, we did not replicate this finding. In addition, recent work suggests that helping without social reward does not engage the NAc (Ruzal et al., 2026). Thus, it’s possible that varying social reward expectations across groups may underly the differences in striatal c-Fos levels seen here.

Within the hypothalamus regions, only the PVN, a classic stress response region (Herman and Cullinan, 1997; Herman and Tasker, 2016), was significantly more active in ELA-conditions. Increased c-Fos levels have consistently been observed in MS-reared animals in stress response regions like the PVN (Fóscolo et al., 2022; Sanders and Anticevic, 2007; Zhang et al., 2012), supporting the general conclusion that ELA sensitizes or primes stress responsive regions. The LHb is another stress-responsive region that was significantly more active in ELA animals on the final day of testing; exposure to a trapped cagemate is presumably aversive, and our findings here fit with prior work suggesting that maternal separation primes the LHb to be hyperactive following a stressful experience (Webster et al., 2023).

### 4.5 Correlations between c-Fos and behavior

Here, helping behavior was significantly correlated with c-Fos levels in several regions of the prefrontal cortex (PFC). In general, the PFC is highly activated in empathy and prosocial based tasks in humans (Bellucci et al., 2020). Here, the prelimbic cortex (PrL) and several regions of the OFC were specifically correlated with prosocial helping behavior. These findings align well with human studies suggesting that more ventral and medial regions of the PFC are associated with prosociality (Bellucci et al., 2020; Lockwood et al., 2024) and are in alignment with prior work in rodents (Ben-Ami Bartal et al., 2021; Breton et al., 2022; Ruzal et al., 2026). The OFC plays a critical role in value-based decision making (Gore et al., 2023; Wallis, 2007) and is involved in cooperation-based tasks in rodents (Jiang et al., 2021), making it a central target for future studies on prosocial behaviors. Correlations were also found between helping behavior and the CA1, which is necessary to acquire and learn socially transmitted information from a peer, and can later guide prosocial choices (Corrêa et al., 2026; La Greca et al., 2026; Walsh et al., 2023). Interestingly, c-Fos levels in the CA1 were not significantly different across ELA and control-reared conditions here, indicating that although it may play a role in prosocial helping, activity in the CA1 alone did not distinguish groups based on early experiences.

### 4.6 Neural network changes following ELA

Brain-wide correlations of c-Fos provide insight into functional connectivity changes in the rodent brain (Terstege and Epp, 2023; Wheeler et al., 2013) and can offer a valuable look into how brain-wide connectivity may shift following early adversity. Here, we see some common hubs emerge in both control and ELA-reared rats – including the lateral septum (LS) – a region previously identified as central in the helping behavior task (Ben-Ami Bartal et al., 2021). Other regions previously identified as part of a ‘prosocial brain network’, including the anterior insula (AI) and medial orbitofrontal cortex (MO), also appeared in both control and ELA-reared rats with high connectivity. As a majority of both control and ELA-reared rats ultimately became helpers, these data lend support for common underlying neural networks involved in prosocial motivation.

Beyond these shared features, network connectivity differed between control and ELA-reared rats, with an overall more cohesive network in control-reared rats, and more distinct clusters in ELA-reared animals. For example, the medial amygdala (MeA) emerged as a central hub for ELA-reared rats, but not for controls. Notably, the mLHb, also appeared as the top central node in ELA-reared animals, connecting canonical prosocial regions with the rest of the brain, including other stress responsive regions like the PVN and BNST.

Altogether, the increased connectivity of stress-networks seen in ELA-reared animals here may be suggestive of heightened stress sensitivity. Yet, whether these functional connectivity changes are causally related to differences in prosocial behavior requires future work and manipulation studies. These data, though descriptive in nature, provide initial insight into brain mechanisms explaining group differences in prosocial behavior between ELA and control reared rats. Future studies can extend findings to specifically address how connectivity changes in individual animals might influence prosocial behavior, using tracing or manipulation-based techniques.

### 4.7 Limitations & Future Directions

Here, rats were tested in a helping behavior paradigm with their trapped cagemate of the same rearing condition. Future studies could test whether increased helping in ELA-reared rats persists across different social contexts, including interactions with unfamiliar conspecifics or animals of differing rearing conditions. Additionally, it will be important to determine whether heightened helping reflects altered signaling from ELA-reared rats (e.g., increased stress cues) or enhanced responsiveness of ELA animals to distressed conspecifics. Another avenue to explore is whether the type of adversity experienced early in life leads to different prosocial outcomes. For example, it is possible that MS, which has more elements of deprivation, may lead to different social outcomes compared to models of resource scarcity, which have more elements of unpredictability and threat (Gallo et al., 2019). These two ELA models lead to varying changes in maternal care (Demaestri et al., 2022; Granata et al., 2021; Orso et al., 2019), and understanding how maternal behavior is linked to social outcomes remains a critical area of ongoing study. Importantly, the helping behavior task used here allows animals to socially interact upon release. Future work can test variants of the task where animals are freed into separate arenas (Bartal et al., 2011; Ruzal et al., 2026), testing whether social interaction and social reward is critical for the ELA-driven behavioral and neural outcomes observed here. The current study also utilized c-Fos as a marker of neural activity, which comes with temporal limitations (Kovács, 2008; Sheng and Greenberg, 1990). Future work could utilize newer methods such as fiber photometry to record neural activity in real-time as animals perform that task.

## 5. Conclusions

Altogether, our findings suggest that ELA may enhance prosocial helping behavior in adult male and female rats, with engagement of different neural networks compared to control-reared animals. Understanding the environments in which prosocial helping arises, and the neural networks involved, is critical for promoting positive social interactions. The results of the present study hold important implications for understanding empathy-related behaviors in humans and how early experiences like adversity may influence social outcomes.

## Supporting information

Supplemental Materials Compiled

## Acknowledgements

We would like to thank Sheta Roy for her help with breeding and animal colony maintenance. We would also like to thank everyone who provided thoughtful comments on this manuscript, including Dr. Maya Rosen. We also thank the Institute for Chemical Imaging of Living Systems (RRID:SCR_022681) at Northeastern University, and Judith Wopereis and the Center for Microscopy and Imaging (CMI) at Smith College for consultation and imaging support.

