## Supplemental Materials Compiled for "Early life adversity enhances prosocial helping towards a cagemate and reshapes brain-wide neural activity and network organization"

**Figures**

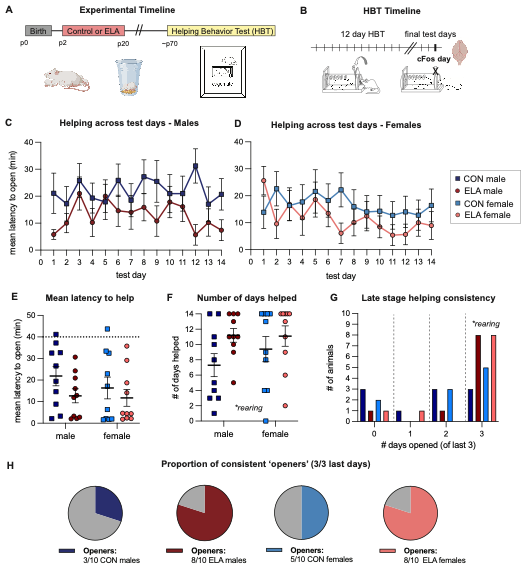

**Figure 1. ELA influenced helping behavior in adult rats.** A) Experimental Timeline. Animals were reared under control (CON) or early life adversity (ELA) conditions from postnatal day (p) 2-p20. All animals started the helping behavior test (HBT) in adulthood. B) Animals underwent 12 days of the HBT followed by a one-week break. There were three final test days; on the last day the door was locked shut such that the free rat could not open it. Brains were collected in a time-locked manner for subsequent analysis. C-D: Helping behavior across the test days in male (C) and female (D) rats. Shown is the average latency to open the door (in minutes) per group. E) The mean latency to open the door averaged across all test days. The dashed line at 40 minutes represents when the door was halfway opened by the experimenter. F) The overall number of days with successful helping by group. G) Late stage helping consistency across the last testing days where animals were able to help (days 12-14). Shown is the number of animals per group who helped on either 1, 2 or 3 of these three testing days. H) The overall proportion of rats that became fully consistent ‘openers’, defined as opening on 3/3 of these last days 12-14. n=10 per group.

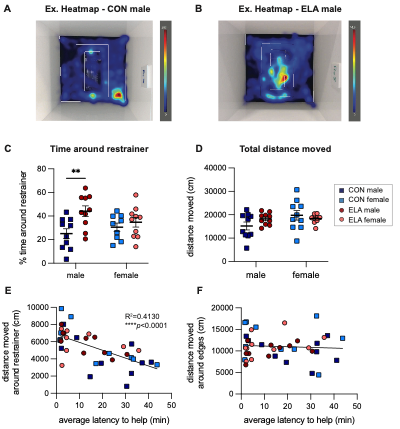

**Figure 2. ELA-reared rats spent more time with their trapped cagemate on the final test day.** (A-B): Representative heatmaps of movement patterns from a control-male (A) and ELA-male (B) on the final test day (day 15). Zones are shown including the zone around the restrainer. C) Percent of time spent in the zone around the restrainer across the 1-hour session. D) Total distance traveled on the final session. (E-F): The distance moved within the zone around the restrainer was significantly correlated with the animal’s latency to help on prior days (E) but was not correlated with distance moved around the edges of the arena (F). n=10 per group.

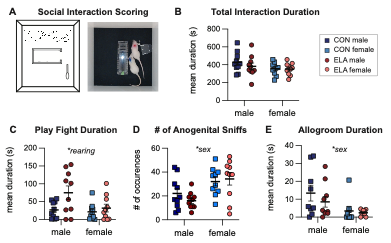

**Figure 3. ELA-reared rats showed increased play fighting upon release of their trapped cagemate.** A) Diagram and representative image of social interaction between two cagemates upon opening the restrainer door. B) The total duration of group social interactions across all social interaction scoring. C) Play fight duration D) The number anogenital sniffs and E) Allogrooming duration. n=10 per group.

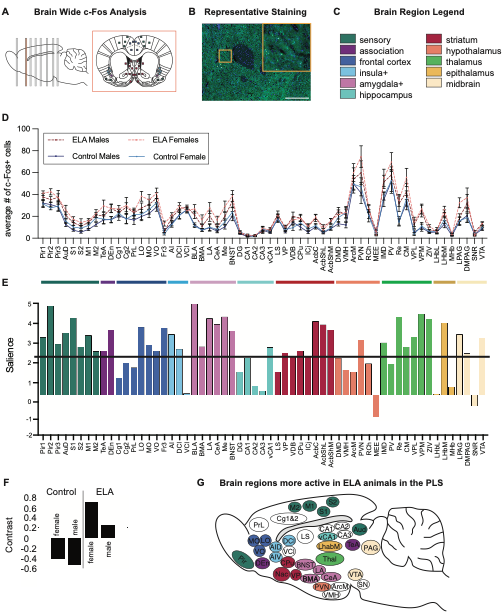

**Figure 4. ELA-reared rats have higher levels of neural activity compared to controls.** A) Diagram showing locations of brain slices used for c-Fos analysis and a representative section from A.P. 1.56. B) Representative c-Fos staining from the nucleus accumbens core. Scale bar is 500μm on the full image, and 50μm on the zoomed in square. C) Legend of brain region categories coded by color, used throughout the manuscript. D) Number of c-Fos+ cells per region (mean ± SEM). E) A salience plot (representing z-scores of bootstrapped tests), with regions crossing the black threshold line significantly (p<0.01) contributing to the contrast shown in (F). The directionality of the bars corresponds with the direction in the contrast graph below. G) A sagittal section with brain regions that crossed the salience threshold in (E) colored.

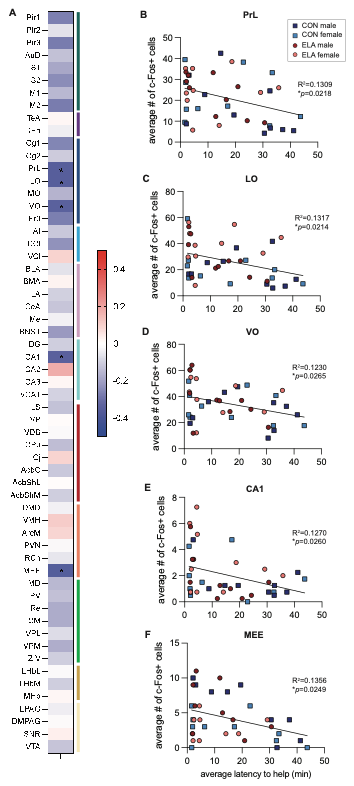

**Figure 5. c-Fos levels in three prefrontal cortex regions were correlated with helping behavior.** A) Heatmap of Pearsons’s correlations (r-values) showing correlations of c-Fos levels in each brain region with the average latency to help during the HBT. Red indicates positive correlations, while blue indicates negative correlations (more c-Fos associated with faster latencies to help). Statistically significant regions are denoted with asterisks. (B-E): Correlation graphs of the five significant brain regions: the PrL (B), LO (C), VO (D), CA1 (E) and MEE (F).

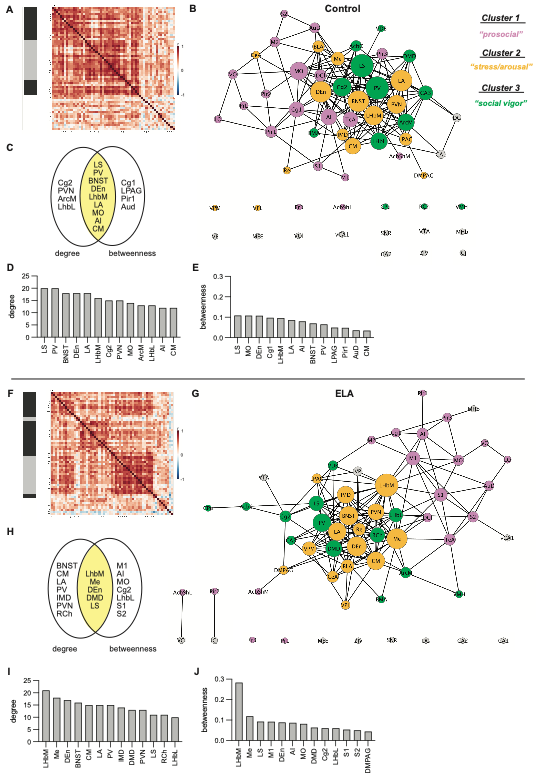

**Figure 6. Different patterns of functional connectivity were observed in ELA compared to control rats.** Correlation matrices (A,D), network maps (B, E) and central hubs (C-E, H-J) for control-reared rats (A-E) and ELA-reared rats (F-J). Networks are color coded based on the three main clusters that emerged from all animals – see supplemental figure S7. The “prosocial” cluster is in pink, the “stress/arousal” cluster is in orange, and the “social vigor” cluster is in green.
